# scTagger: Fast and accurate matching of cellular barcodes across short- and long-reads of single-cell RNA-seq experiments

**DOI:** 10.1101/2022.04.21.489097

**Authors:** Ghazal Ebrahimi, Baraa Orabi, Meghan Robinson, Cedric Chauve, Ryan Flannigan, Faraz Hach

## Abstract

Single-cell RNA sequencing allows for characterizing the gene expression landscape at the cell type level. However, because of its use of short-reads, it is severely limited at detecting full-length features of transcripts such as alternative splicing. New library preparation techniques attempt to extend single-cell sequencing by utilizing both long-and short-reads. These techniques split the library material, after it is tagged with cellular barcodes, into two pools: one for short-read sequencing and one for long-read sequencing. However, the challenge of utilizing these techniques is that they require matching the cellular barcodes sequenced by the erroneous long-reads to the cellular barcodes detected by the short-reads. To overcome this challenge, we introduce scTagger, a computational method to match cellular barcodes data from long-and short-reads. We tested scTagger against another state-of-the-art tool on both real and simulated datasets and we demonstrate that scTagger has both significantly better accuracy and time efficiency.

## 1 Introduction

Single-cell RNA sequencing (scRNA-seq) has advanced biological research to move beyond the macro view provided by bulk RNA sequencing. Technical advances have enabled gene expression analysis at the single-cell level, which has resulted in a greater understanding of cellular diversity in gene expression [1]. scRNA-seq achieves this cell-level resolution by deploying short artificial DNA fragments known as cellular barcodes to tag the RNA fragments in each cell with a unique sequence of nucleotides. Once tagged, complementary DNA strands are created for each RNA strand and are subsequently fragmented, amplified, and loaded onto the sequencing machine to be sequenced in bulk. Following this, the sequencing outputs are demultiplexed, and reads are assigned to different cells *in silico* using the sequenced cellular barcodes. The genes and their RNA expression levels that are attributed to each cell allows the user to classify cells into clusters of cell types or populations that are biologically relevant [2, 3]. Observing the gene expression at the scale of thousands of cells from a single sample is made readily accessible using droplet-based technologies such as the 10x Genomics Chromium approach [4].

While gene expression is valuable to reveal biological processes, it tells only a part of the transcription story. Genes are comprised of various isoforms that differ in the order and composition of their respective exons, which have significant biological implications in subsequent protein translation and function. Unfortunately, evaluating these splicing variations in RNA isoforms is limited and often impossible when using standard scRNA-seq techniques, such as the 10x Genomics Chromium protocol, since they rely on short-read (SR) sequencing platforms, often Illumina where 150 total base pairs are typically sequenced at a time. The reason SR sequencing is utilized by scRNA-seq is due to its low error rate (∼ 0.1% [5]) which allows for easy demultiplexing of cellular barcodes. Additionally, SR platforms relative affordability allows for high coverage sequencing that enables the estimation of gene expression. However, because of this reliance on SR sequencing, scRNA-seq fails to capture the full picture of transcripts that are much longer than what the current SR platforms can generate.

Long-read (LR) sequencing technologies, such as Oxford Nanopore Technologies (ONT), have demonstrated their capacity to identify full-length isoforms, albeit using bulk RNA that is not multiplexed at the cellular level [6, 7]. Recently, new and promising library preparation techniques aim to take advantage of both LR RNA sequencing and SR single-cell RNA sequencing [8, 9, 10]. These techniques typically modify existing SR scRNA-seq protocols by splitting the RNA material, after it is tagged with the cellular barcodes, into two pools: i) one for Illumina SR sequencing and ii) one for LR sequencing. The high-level overview of this library preparation technique is illustrated in Figure 1.

**Figure 1:**
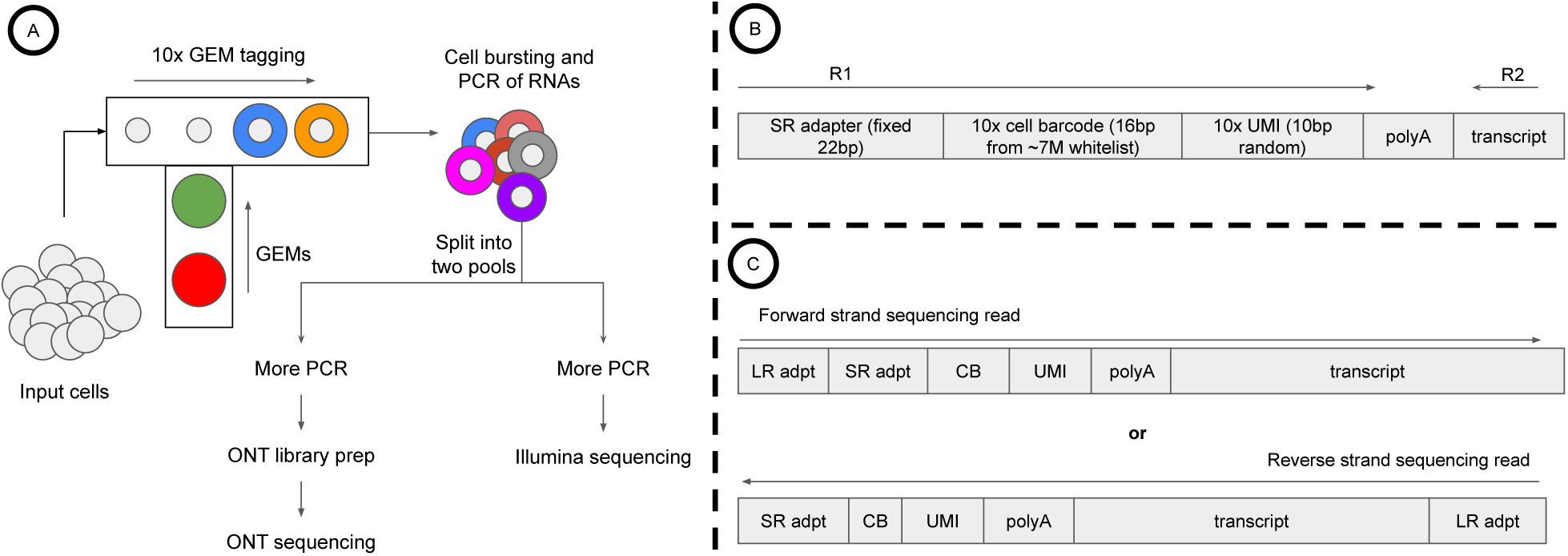
**A)** Overview of the library preparation: Microfluidic chips are used to generate 10x Chromium GEMs which tag the RNA transcripts with cellular barcodes. After the transcripts are tagged, the GEMs are burst, and the tagged RNA material is split into two pools of sequencing, one for SRs and one for LRs. **B)** The SR template. Inside the GEMs, RNA transcripts are tagged with an Illumina adapter that is a fixed sequence, followed by a 16bp cellular barcode (CB), followed by a random 10bp sequence for the unique molecular identifier (UMI). **C)** The LR template. Note that the LR template is essentially the same as the SR template with the LR sequencing adapter added. Depending on the specifics of the library preparation, LRs may sequence the forward or the reverse strand of the RNA molecule. In either case, we expect the cellular barcode to be adjacent to the SR adapter sequence.

These techniques are designed to enable the clustering of cells into types, using cell-specific gene expression, while taking advantage of the LR data to identify alternatively spliced isoforms and map them to cells (and subsequently into cell types) using the shared cellular barcodes. However, this mapping process is computationally not trivial because of the level of noise exhibited by LR sequencing platforms (10% − 20% [11]), which frequently introduces errors to the cellular barcodes sequences. Current methods to associate LRs to cellular barcodes include SiCeLoRe [12] and FLAMES [9]: SiCeLoRe relies on the exact matching of permutated mutations of the LR barcode sequences, while FLAMES relies on the alignment of the barcodes to the LRs using dynamic programming. Such approaches are theoretically computationally prohibitive for a large number of cells and reads.

Therefore, we present scTagger, a novel computational method for efficiently and accurately identifying and matching single-cell barcodes in split hybrid LR-SR scRNA sequencing experiments that can scale for a large number of reads and cells. scTagger uses a trie-based data structure to efficiently match the identified barcodes in the SRs to the LRs while allowing for non-zero edit distance matching. Using both real and simulated datasets, we show that scTagger has accuracy on par with an exact but computationally intensive dynamic programming-based matching approach while being orders of magnitude faster. Furthermore, we show that scTagger has much higher accuracy and is much more time-efficient than other available tools.

## 2 Results

### 2.1 scTagger overview

The hybrid SR-LR protocol we refer to in the Introduction starts with RNA from single cells of a given sample. The RNA molecules of each cell barcoded with a fixed-length barcode^1^ sampled from a pool of over six million (10x Chromium v3 Chemistry) possible whitelisted 16bp barcodes. This RNA material is split into two pools, one pool for SR Illumina sequencing, which is used to produce a gene expression matrix, and the other pool for LR ONT sequencing, which is used to identify isoforms. The expected template sequence for the SRs and the LRs is illustrated in Figure 1B and 1C.

scTagger aims to match each LR to one of the cellular barcodes that tagged the input cells. To achieve this goal, scTagger proceeds in three stages:

i. Pre-processing the SRs and identifying the barcode of each SR. In this stage, scTagger selects a *small* subset (proportional to the number of input cells) of barcodes that cover most of the SRs using the frequencies of the barcodes appearing in the SRs. The small size of this subset allows for efficient matching without significant loss of data.
ii. Locating a short segment on each LR where the barcode is expected to be present. In this stage, scTagger exploits the apriori knowledge about the template of the LRs (Figure 1C) and uses the alignment of the fixed Illumina adapter sequence to each of the LRs to identify the segment where the barcode is present.
iii. Matching the SR barcodes to the LR segments. scTagger achieves this matching by using a trie data structure [13]. The trie is modified to allow matching each LR segment to the barcode that requires the least number of edits (e.g. substitutions, deletions, or insertions) to form an exact match with the segment. The maximum number of allowed edits for a barcode-segment match is set by the user and is limited by default to 2 errors.

Figure 2 presents an schematic overview of these stages and the STAR Methods section provides details of them.

**Figure 2:**
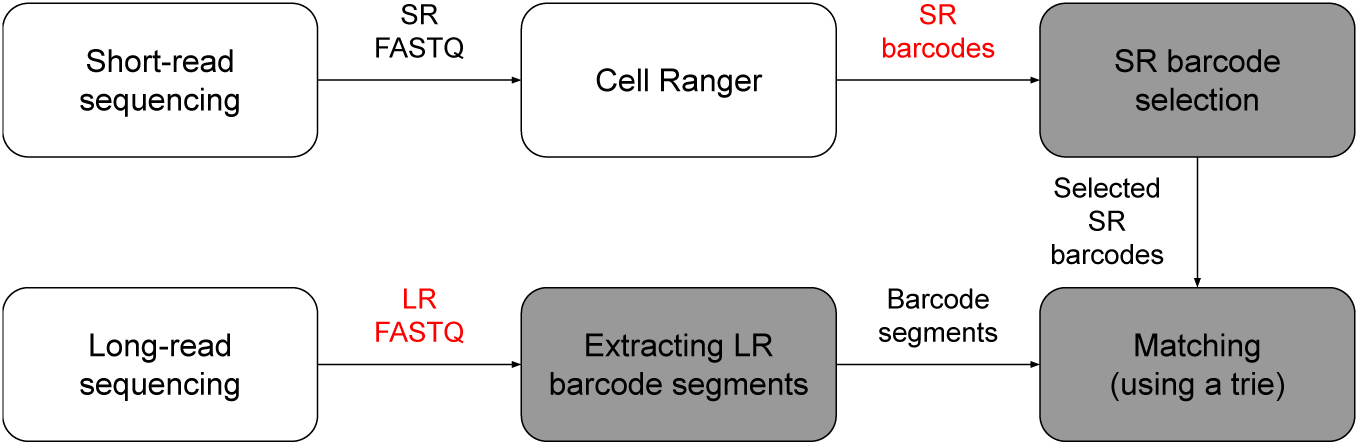
The three stages of scTagger are shaded while the data inputs are coloured in red.

### 2.2 Experimental setup

We assessed the tested methods, FLAMES (GitHub commit 18fb83c) and scTagger, on both real and simulated datasets. Note that we attempted to run SiCelore but failed to run it successfully due to runtime errorswhich is concordant with the reporting from FLAME’s paper on SiCelore [9]. In addition to FLAMES and scTagger, we also ran a baseline brute-force method of matching the cellular barcodes in the LRs by performing a *glocal* dynamic programming based alignment of all the selected barcodes (stage 1) to all the extracted LR segments (stage 2). We will refer to this baseline method as the Brute-force matching method in this manuscript. Note that all the matches found by scTagger exactly mirror those found by the Brute-force method as long as they have a maximum edit distance of e = 2, the maximum allowed distance set in scTagger. Additionally, according to FLAMES’ manual, we are supposed to run FLAMES with the full 10x Genomics barcode whitelist, which includes over 6 million barcodes. However, doing so resulted in FLAMES not terminating even after running for 24 hours. Therefore, we modified the way we run FLAMES by feeding it the selected subset of barcodes found by scTagger (stage 1). Thus, the results we present here for FLAMES depend on some of the output of our own method.

#### 2.2.1 Dataset generation

We generated three real and three simulated datasets. The details of the generating procedures for both follows below, and the summary statistics for each dataset are detailed in Table 1.

**Table 1:**
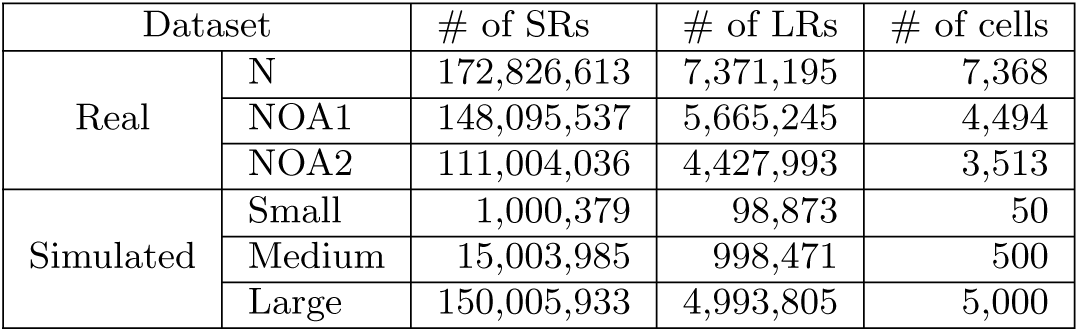
Summary of used datasets. The number of cells for real datasets is estimated by 10x Genomics Cell Ranger v3.0.2.

| Dataset |  | # of SRs | # of LRs | # of cells |
| --- | --- | --- | --- | --- |
| Real | N | 172,826,613 | 7,371,195 | 7,368 |
|  | NOA1 | 148,095,537 | 5,665,245 | 4,494 |
|  | NOA2 | 111,004,036 | 4,427,993 | 3,513 |
| Simulated | Small | 1,000,379 | 98,873 | 50 |
|  | Medium | 15,003,985 | 998,471 | 500 |
|  | Large | 150,005,933 | 4,993,805 | 5,000 |

##### Real datasets

We isolated cells from three different male infertility patients. The three samples were extracted from the testis tissue using the protocol developed by Valli et al. [14]. The first sample was extracted from a normal human testis biopsy, while the second and third samples were extracted from human testis biopsies diagnosed with non-obstructive azoospermia, a severe abnormality in spermatogenesis resulting in failure to produce sperm. We name these three samples N, NOA1, and NOA2, respectively. Libraries were prepared in accordance with the protocol developed by Gupta et al. [10] for 10x Genomics Chromium Single Cell 3’ v3.0 and Oxford Nanopore Technologies PromethION platform. For the cDNA amplification step, we applied 17 cycles of PCR. Then, the amplified and barcode tagged cDNA material was split into two pools. The first pool was sequenced on the Illumina HiSeq platform with a read length of 2×150bp. The second pool was the input for the ONT cDNA sequencing protocol, for which we used the SQK-PBK004ONT ONT kit. For this pool, end repair, A-tailing, and ligation steps were performed, and 11 cycles of PCR were applied using a starting cDNA amount of 50ng.

##### Simulated datasets

In addition to the real datasets, it is important to assess the accuracy of the different tested methods on simulated datasets for which we can obtain the ground truth of barcode matching between the LRs and SRs. Therefore, we built a simulation pipeline to generate SR and LR data with known cellular barcode matching. The structure of the simulation pipeline is outlined in Figure 3. Note that we use both Minnow [15] and Badread [16] simulators as components of our simulation pipeline. In total, we generated three simulated datasets. As part of the input for simulating all three datasets, we used the transcriptome mapping of the SRs of the real sample N as a basis to simulate the differential expression of gene isoforms according to Minnow’s instructions.

**Figure 3:**
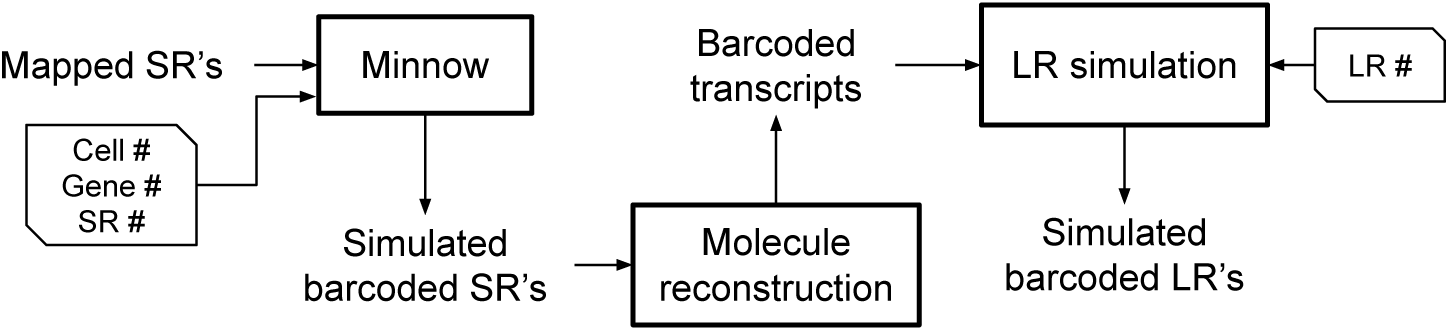
Overview of simulation pipeline: The SRs mapping to the transcriptome reference is used by Minnow to model differential isoform expression within each gene. The number of genes, SRs, and cells to be simulated are passed to Minnow. We then reconstruct the transcript molecule (including SR adapter, barcode, and polyA tail) that each simulated SR was generated from. These reconstructed barcoded transcripts are then passed to the LR simulation module which wraps around Badread simulator to generate the LRs. Note that the full simulation pipeline is available on our GitHub repository.

### 2.3 Barcode selection and SR coverage

The first stage of scTagger filters and selects SR barcodes. Here, we need to examine how well the selected barcodes cover the SR data and how little value adding more barcodes brings. A significant amount of the SR coverage is wasted on barcodes that did not tag any input cells in the library preparation and are therefore not useful for downstream analysis.

In Figure 4, we plot the proportion of SRs that are covered by retaining each additional 1,000 barcodes in order of the most-to-least frequent. As we observe in the figure, the contribution of adding each additional 1,000 barcodes diminishes quickly. Therefore, we observe that our barcode selection method (with a threshold of 0.5%) achieve its conservative goal while producing a relatively small set of error-corrected barcodes. The exact number of selected SR barcodes and the proportion of SRs they cover is detailed in Table 2. Due to this barcode selection process, we lose between 9% and 17% of the SR throughput. These selected barcodes are the ones used for matching in all the tested methods. Note that in all three cases, the number of selected barcodes exceeds the expected number of cells in each sample (Table 1).

**Figure 4:**
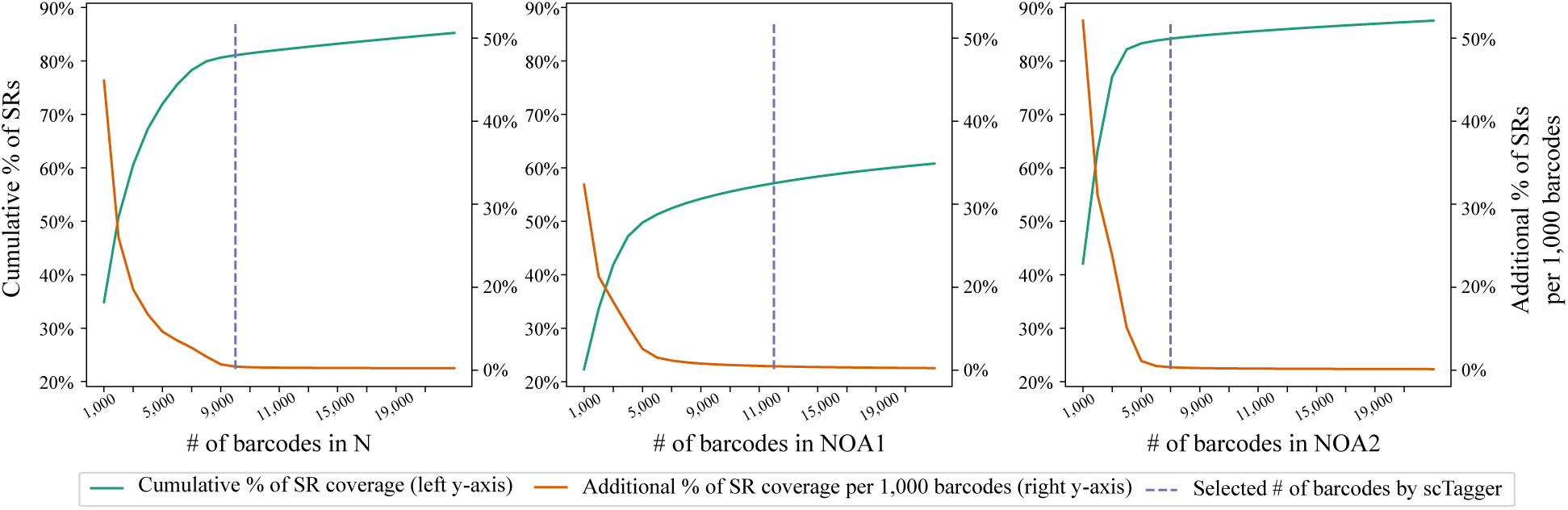
Cumulative SR coverage with batches of 1,000 barcodes for the real datasets.

**Table 2:** The number of unique barcodes in each real dataset and the percentage of SRs that they cover. Here we compare the set of all barcodes we observe in the SRs vs the set of selected barcodes by scTagger. Note that some SRs do not have any observed barcodes.

| Dataset | All barcodes |  | Selected barcodes |  | % of SRs lost due to barcode selection |
| --- | --- | --- | --- | --- | --- |
|  | # of unique barcodes | % of SRs covered | # of unique barcodes | % of SRs covered |  |
| N | 1,723,938 | 97% | 10,000 | 80% | 17% |
| NOA1 | 1,063,678 | 72% | 14,000 | 55% | 17% |
| NOA2 | 859,034 | 94% | 7,000 | 85% | 9% |

### 2.4 LR coverage and segment extraction

The second stage of scTagger extracts LRs barcode segments. scTagger uses the alignment of the Illumina sequencing adapter to the LRs to identify the segment containing the cellular barcode in each LR. In Figure 5, we plot the histogram of the LRs by the edit distance of the successful adapter alignments. We observe that we are able to find successful adapter alignments in over 75% of LRs across all real datasets. For the simulated datasets, we observe that scTagger is able to find successful adapter alignments on ∼ 92 − 93% of the LRs.

**Figure 5:**
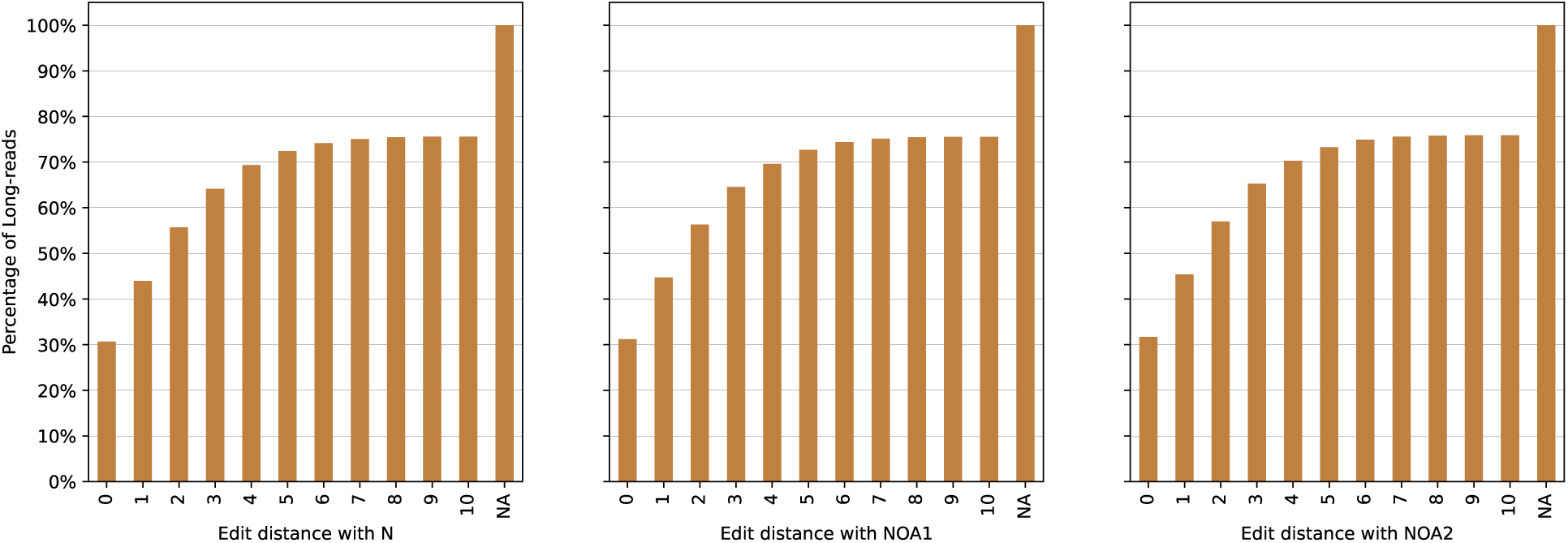
Cumulative percentage of LRs distance with a successful SR adapter alignment. NA refers to LRs with no alignment *≤* 10 edits.

### 2.5 Accuracy of LR barcode matching

The final and main stage of scTagger is matching the selected SR barcodes with the extracted LR segments. Here, we use the simulated datasets to assess the accuracy of the different methods. For every tested method, we label each simulated LR as: (i) a match if the method outputs only the one barcode we simulated for the LR; (ii) an ambiguous match if the method outputs multiple barcodes and one of them is the simulated barcode; (iii) a mismatch if none of the output barcodes is the simulated barcode; and finally (iv) a skip if the method does not output any barcodes for the LR (due to the edit distance limit or due to not finding any feasible adapter alignment in the second stage of scTagger). Note that while ambiguous matches include the correct barcode, they are not readily useful for any downstream analysis. Table 3 details the accuracy statistics for all the tested methods.

**Table 3:** Accuracy on simulated datasets.

| Dataset | Method | Matches | Ambiguous matches | Mismatches | Skipped |
| --- | --- | --- | --- | --- | --- |
| Small | scTagger | 47.68% | 9.98% | 4.57% | 37.76% |
|  | Brute-force | 53.22% | 23.82% | 14.77% | 8.19% |
|  | Flames | 27.40% | 0.00% | 15.92% | 56.68% |
| Medium | scTagger | 51.60% | 7.10% | 4.54% | 36.76% |
|  | Brute-force | 55.55% | 20.03% | 17.37% | 7.05% |
|  | Flames | 28.37% | 0.00% | 22.10% | 49.52% |
| Large | scTagger | 53.26% | 6.24% | 5.87% | 34.62% |
|  | Brute-force | 54.91% | 18.42% | 19.62% | 7.04% |
|  | Flames | 30.05% | 0.00% | 20.34% | 49.61% |

As we observe in Table 3, scTagger significantly outperforms FLAMES in its accuracy. And despite the low edit distance limit we use in scTagger (e = 2), scTagger has a very similar match rate to the Brute-force baseline method. As a matter of fact, we observe that allowing for higher edit distance increases the mismatch and ambiguous match rates.

While we cannot establish a ground truth to calculate the accuracy of scTagger in matching LRs of the real datasets, we can still compare scTagger matching to the baseline Brute-force method. Note that the differentiating parameter between these two methods is the choice of the edit distance limit in scTagger; if we set that parameter to the barcode size, the two methods produce identical results. Figure 6 depicts the distribution of the real datasets LRs by whether they are: (i) skipped; (ii) matched with edit distance ≤ 2; (iii) matched with edit distance > 2; and (iv) matched by a single unique barcode. As we observe in these UpSet [17] plots, only 0.8% to 2.2% of all the LRs have unambiguous matches and yet are lost by scTagger because of having an edit distance limit of 2 edits. Thus, the loss of matches due to setting the error limit to 2 is very small in practice. The remaining ambiguous matches, i.e. those with more than one barcode match, are not readily usable by downstream analyses. Additionally, when comparing FLAMES and scTagger matches with the Brute-force method, we observed that the percentage of the LRs that are matched by FLAMES and the Brute-force methods but not scTagger is negligible (∼ 0.1%). In contrast, the percentage of the LRs that are matched by scTagger and the Brute-force methods but not FLAMES is very significant (∼ 26% to ∼ 28%). Table S1 of the Supplementary Material presents the statistics of comparing the three methods’ matches on the real datasets.

**Figure 6:**
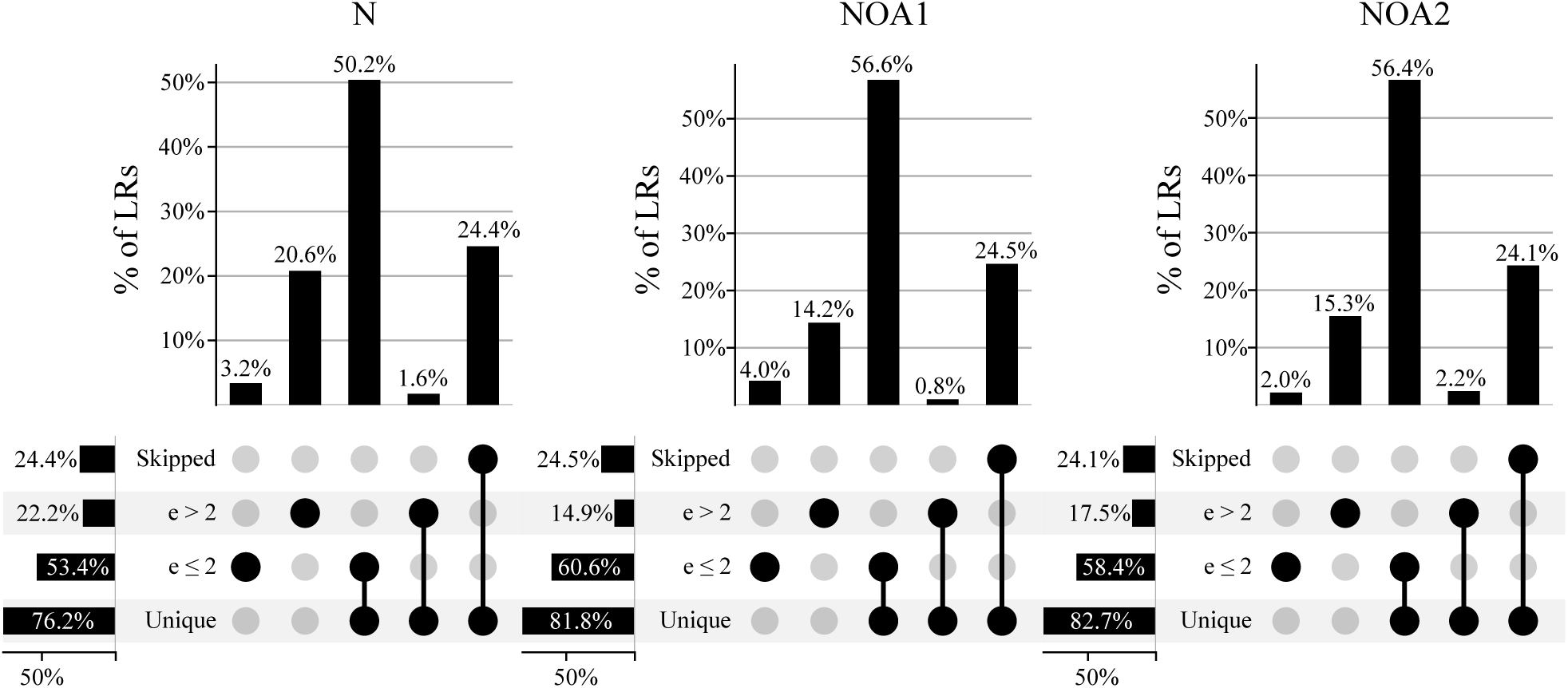
Distribution of matches in the real datasets according to the Brute-force matching method.

### 2.6 Runtime and memory

Table 4 shows the memory usage and runtime of the different stages of scTagger as well as FLAMES and the Brute-force method. As we observe in Table 4, scTagger matching stage has the smallest runtime footprint compared to the other methods. Unlike FLAMES, scTagger can take advantage of multiple CPU cores and completes computing in orders of magnitude less time than FLAMES or the Brute-force method. The only computational resource that scTagger lags behind the other tested tools is memory usage. However, its memory usage for these real datasets is well within the acceptable limits of modern personal computers.

**Table 4:**
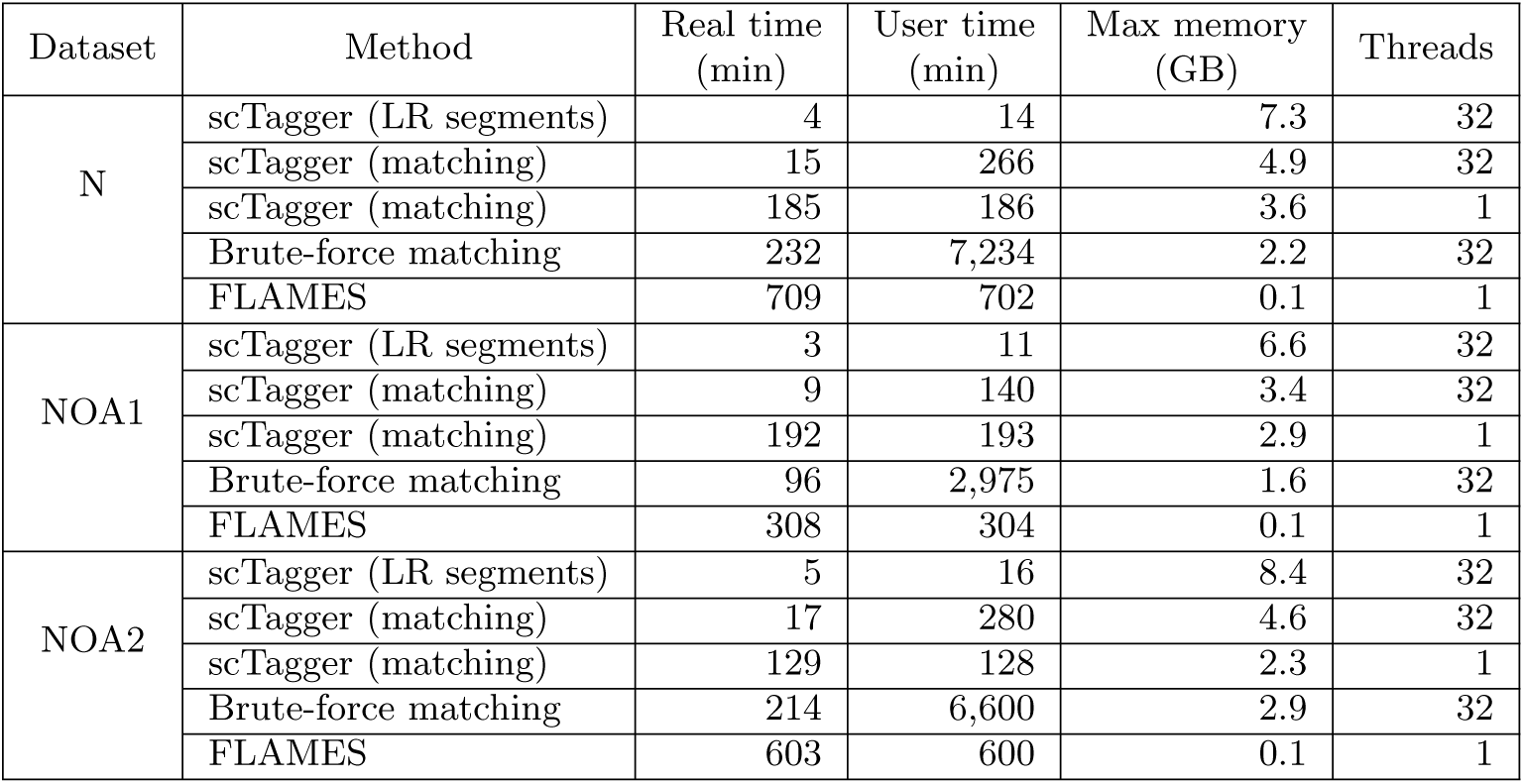
Time and memory usage on the real datasets as reported by GNU Time.

| Dataset | Method | Real time<br>(min) | User time<br>(min) | Max memory<br>(GB) | Threads |
| --- | --- | --- | --- | --- | --- |
| N | scTagger (LR segments) | 4 | 14 | 7.3 | 32 |
|  | scTagger (matching) | 15 | 266 | 4.9 | 32 |
|  | scTagger (matching) | 185 | 186 | 3.6 | 1 |
|  | Brute-force matching | 232 | 7,234 | 2.2 | 32 |
|  | FLAMES | 709 | 702 | 0.1 | 1 |
| NOA1 | scTagger (LR segments) | 3 | 11 | 6.6 | 32 |
|  | scTagger (matching) | 9 | 140 | 3.4 | 32 |
|  | scTagger (matching) | 192 | 193 | 2.9 | 1 |
|  | Brute-force matching | 96 | 2,975 | 1.6 | 32 |
|  | FLAMES | 308 | 304 | 0.1 | 1 |
| NOA2 | scTagger (LR segments) | 5 | 16 | 8.4 | 32 |
|  | scTagger (matching) | 17 | 280 | 4.6 | 32 |
|  | scTagger (matching) | 129 | 128 | 2.3 | 1 |
|  | Brute-force matching | 214 | 6,600 | 2.9 | 32 |
|  | FLAMES | 603 | 600 | 0.1 | 1 |

## 3 Discussion

We presented a novel computational method, scTagger, to match SR data to LR data using their cellular barcodes. We show that scTagger achieves this goal more efficiently and accurately than the other alternative methods. In the future, we plan to extend scTagger method to handle matching of the unique molecular identifier sequences present in the SR and LR data. We also plan to improve on scTagger’s memory use by streamlining how it reads and processes the input.

### 3.1 Limitation of the study

The main limitation of the current form of scTagger is that it is specific for 10x Genomics scRNA-seq. Note that is possible to readily extent support for other scRNA-seq platforms if their resulting LR template includes some fixed sequence next to the barcode. Additionally, scTagger takes advantage of the fact that the number of cells (and thus the number of relevant barcodes) is on the manageable order magnitude of thousands to tens of thousands. Therefore, we expect scTagger in its current design not to scale well if the number of relevant barcodes becomes much bigger than that.

## 4 Acknowledgements

## Funding

This research is funded in part by the National Science and Engineering Council of Canada (NSERC) CREATE Training Program in High-Dimensional Bioinformatics Scholarship to G.E.; NSERC Alexander Graham Bell Canada Graduate Scholarship-Doctoral (CGS D) to B.O.; NSERC Discovery Grants to F.H (RGPIN-05952) and C.C (RGPIN-03986); the Michael Smith Foundation for Health Research (MSFHR) Scholar Award to F.H. (SCH-2020-0370); and Canadian Institute for Health Research (CIHR) early career investigator grant (161357) to R.F.

We would like to thank Reza Soltani for his thoughtful input into the development and calculating the time complexity of the trie data structure in scTagger and Fatih Karaoğlanoğlu for his feedback on designing the simulation pipeline.

## 5 Author contributions

Conceptualization: G.E., B.O., R.F. and F.H.; Data curation: G.E., B.O., M.R., R.F. and F.H.; Formal analysis: G.E. and B.O.; Funding acquisition: G.E., B.O., C.C., R.F. and F.H.; Investigation: G.E., B.O. and F.H.; Methodology: G.E., B.O. and F.H.; Resources: R.F. and F.H.; Software: G.E. and B.O.; Supervision: C.C., R.F. and F.H.; Validation: G.E. and B.O.; Visualization: G.E. and B.O.; Writing – original draft: G.E., B.O. and F.H.; Writing - review & editing: G.E., B.O., M.R., C.C., R.F. and F.H.;

## 6 Declaration of interests

The authors declare no competing interests.

## STAR ⋆ Methods

### Algorithmic details of scTagger stages

#### SR barcode selection stage

In the first stage, scTagger uses 10x Genomics Cell Ranger [4] that performs genomic mapping, filtering, barcode counting, and UMI counting to build a map from the SRs to their error-corrected cellular barcodes. Ideally, the set of error-corrected barcodes in this map should have the same size as the number of cells used in the library preparation. However, the number of barcodes in this map ends up being orders of magnitude larger than the number of input cells. We believe this is due to library preparation artifacts and sequencing errors. Therefore, it is important to filter out these noisy barcodes before matching them to the LRs. 10x Genomics library protocol draws barcodes from an original whitelist of more than six million barcodes. In our experiments, typically, we observe over a million unique error-corrected barcodes in the SRs (see Table 2 for details). The vast majority of these error-corrected barcodes have very low frequencies, appearing only in a few SRs. These low-frequency barcodes cannot be utilized by downstream scRNA-seq analysis tools. Therefore, we can select a drastically smaller subset of the 10x Genomics barcode whitelist, that will nevertheless cover a majority of SRs, by filtering out the lowest-frequency barcodes.

We achieve this by sorting the set of error-corrected barcodes that are detected by Cell Ranger by the frequency of which they appear in the SRs, in descending order. We then examine the sorted barcodes iteratively in increments of 1,000 barcodes at a time; if the sum of SRs retained as a result of retaining the current 1,000 barcode increment represents less than t% (by default t = 0.5) of the total number of SRs, we stop retaining barcodes. Otherwise, we proceed to the next 1,000 barcode increment. In practice, this process typically reduces the number of barcodes from about a million barcodes to less than a few tens of thousands of barcodes which is higher than the number of input cells. At the same time, this process retains up to 70% of all the SRs. The small size of the selected barcode set allows us to match the LRs to the barcodes much more efficiently than if we were to include the many low-frequency barcodes which are not useful for downstream analyses.

#### Extracting LR barcode segments stage

From the library preparation and its resulting LR template (Figure 1C), we expect the cellular barcodes to be located within specific narrow ranges on the LRs. Thus, if we encounter an adapter alignment outside these expected ranges, we can suspect that it is an erroneous alignment. We believe it is critical to identify these ranges from the sample data directly, rather than rely on theoretical preset ranges. Detecting these possible ranges from the sample’s data allows scTagger to be robust against library preparation artifacts or modifications that may affect the structure of the LR template. We can locate the adapter on the LRs by performing a *glocal* alignment of the adapter to the LRs (i.e., performing a global alignment that does not penalize prefix and suffix deletions on the target). This takes *O*(*R*×*a*) time, where *R* is the total length of the LRs and *a* is a small constant equal to the length of the adapter string^2^. To perform this alignment, we use the edlib library [18]. In Figure 7, we plot the distribution of mapping locations that are used by scTagger to detect the ranges on a real dataset (dataset details in the Results section). As we can observe, the adapter alignment has distinct peaks on each of the forward and reverse strands of the LR data. scTagger automatically detects the ranges around these peaks and uses these detected ranges to reject any adapter alignments that fall outside. This detection process is detailed in Algorithm 1 of the Supplementary Material. Finally, for each successful adapter alignment scTagger extracts the 20bp LR segment starting from the end of the adapter alignment (16bp for the barcode length and an extra 4bp to allow for a small error margin).

**Figure 7:**
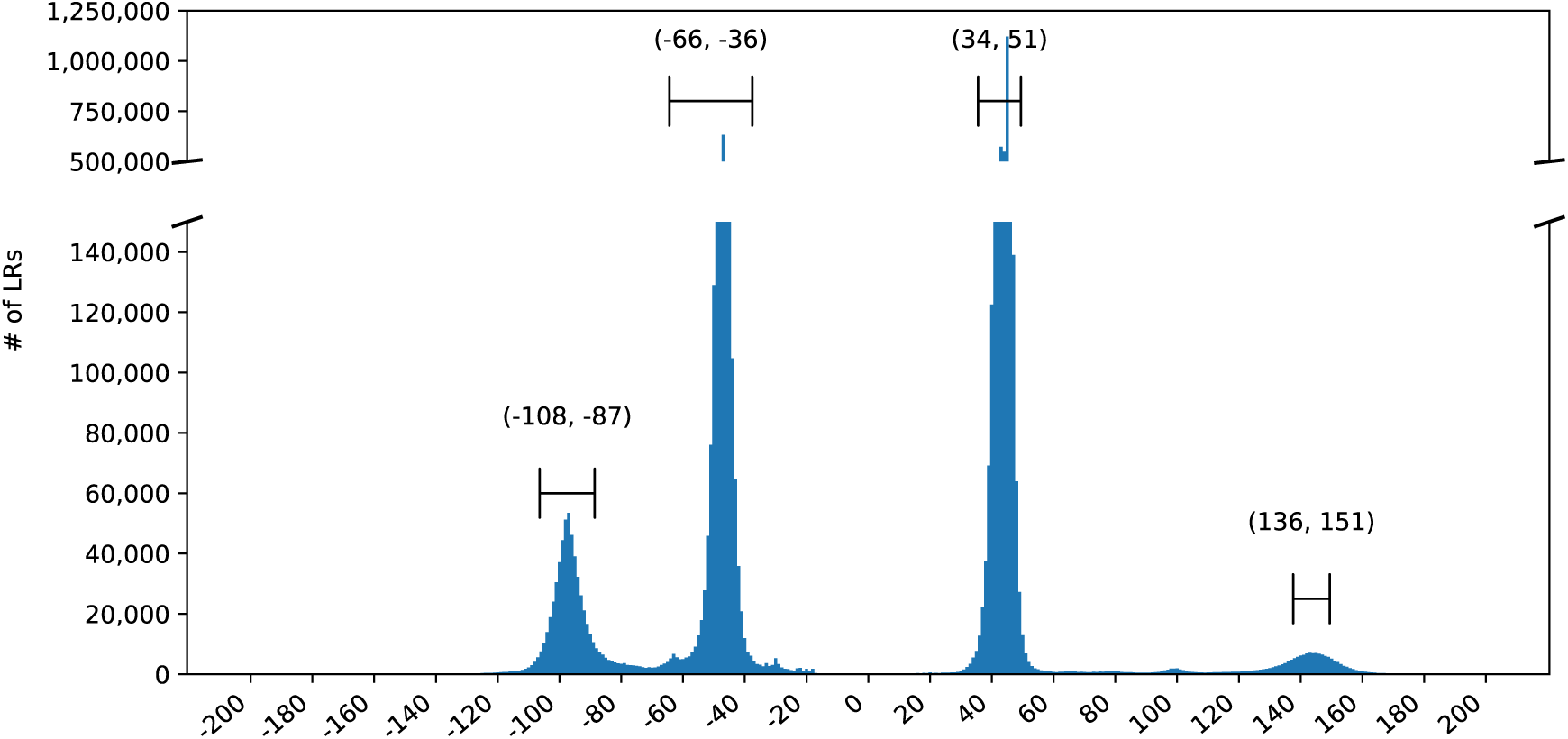
Optimal alignment locations of the SR adapter to the LRs. Negative locations are on the reverse strand. Automatically selected ranges around the peaks for sample N are indicated on the plot.

#### Matching LRs and selected barcodes stage

In this stage, scTagger matches the extracted LRs segments to the selected subset of SR barcodes by finding the best barcode alignment to each LR segment. We use a trie structure to store all k-mers and use SR barcodes as a query. Using a trie allows us to find all LR matches for a given barcode in one query. Also we use a simple unit-cost Levenshtein edit distance since it is more efficient to compute than other more complex edit distances (e.g., affine gap) and should be sufficient for the short length of the barcodes. Computing the minimum (Levenshtein) edit distance between all considered barcodes and LRs by dynamic programming results in a time complexity that is too large: *O*((*N* × *S*) × (*M* × *L*)), where *N* is the number of LRs, *S* is the size of each LR segment, *M* is the number of barcodes, and *L* is the length of each barcode. Instead of using such a brute-force approach, we opted to use a trie data structure to compute the optimal edit distances between SR barcodes and the LR segments up to an assumed maximum edit distance of *e*. We begin by constructing the trie by inserting the LR segments into the trie. Then, we query the trie with each barcode in a fashion that allows for edit errors (insertions, deletions, and mismatches). Note that we use the Numpy [19] library for our implementation of the trie.

#### Trie construction

We extract all the *k*-mers from the LR barcode segments and insert them into the trie. To allow up to *e* edits in matching the barcodes to the LR segments, we choose 16 +*e* as the *k*-mer size (16 for the barcode size plus *e* possible insertion errors). This trie construction step is similar to the classical insertion in tries. Additionally, we maintain a map of all the trie nodes at tree depths ≥ 16 − *e*; any barcode query terminating in shallower nodes will result in alignments errors *> e*. Each node in the map will be associated with the set of LRs with *k*-mers that threaded through the node during their insertion into the trie. This will allow us to find all barcode alignments to LR barcode segments with up to *e* deletions, insertions, or mismatches. Figure S1 in the Supplementary Material illustrates an example of the trie construction process.

#### Querying the trie

For each barcode, we query the trie by searching it recursively, starting from the root. Each recursive call includes the information on how many characters we matched so far on the barcode and how many errors we are still allowed to have. For deletion edits, the recursive call stays on the same node while incrementing the query’s index. For insertion edits, the recursive calls would advance to each of the children of the current node while maintaining the current query index. For mismatch edits, we go to any children node that is not matched by the character at the query index. The recursive calls terminate when the error allowance is negative or the length of the barcode is exhausted. The result of this search is the set of LR barcode segments that contains *k*-mers that are at most *e* edits away from the query barcode. In the Supplementary Material, Algorithm 2 presents the pseudocode of querying the trie and Figure S2 illustrates how we query the trie with a barcode while allowing for these edit errors.

The time complexity of this stage is defined in two parts. First, the trie construction step takes *O*(*N* × *S* × *k*) time where *N* is the number of LRs, *S* is the size of each LR segment, and *k* is the *k*-mer size. This is practically linear since *S* and *k* are small constants. Second, for the querying step, the time complexity is *O*(*M* × 4^*e*^ × (*L* + *e*)^*e*+1^), where *M* is the number of barcodes, *L* is the size of each barcode, and *e* is the maximum allowed edit distance. Details of calculating this upper bound on the time complexity are available in Theorem 6 in the Supplementary Material. Later in the Results section, we show that the vast majority of LRs with *>* 2 edits have no unique barcode match and are therefore not useful for downstream analysis. Therefore, by default, we set *e* = 2. This maintains a runtime that is faster than the brute-force alignment-based method’s runtime while sacrificing little in terms of results.

#### Parallelizing trie construction and querying

To enable parallelized computation of the trie, we partition the space of all possible *k*-mers into disjoint sets. Each available CPU thread can then process the *k*-mers belonging to each partition independently by constructing a trie using the partition’s *k*-mers and querying the barcodes on the constructed trie. The main thread can then collate the query results from all the threads, discarding any sub-optimal barcode-LR matches.

In scTagger, we partition the *k*-mers by their prefix of length *p*. Thus, the number of partitions is 4^*p*^ since the nucleotide alphabet size is 4. We set *p* such that the number of partitions is at least equal to the number of available CPU cores. This partitioning scheme allows for simple data parallelism while resulting in a similar CPU-bound time complexity as with the single-thread (single-trie) implementation discussed above. This is because the only major overhead here is the collation of the results from the threads by the main thread.

## Supplementary Material

**Table S1:** Statistics comparing the matches of FLAMES and scTagger against the Brute-force method.

| Sample | Brute matched? | Brute match unique? | scTagger = Brute? | FLAMES = Brute? | scTagger = FLAMES? | % of LRs | # of LRs |
| --- | --- | --- | --- | --- | --- | --- | --- |
| N | ✓ | ✓ | ✓ | ✓ | ✓ | 23.5% | 1,734,479 |
|  | ✓ | ✓ | ✓ | ✗ | ✗ | 26.7% | 1,965,586 |
|  | ✓ | ✓ | ✗ | ✓ | ✗ | 0.1% | 3,742 |
|  | ✓ | ✓ | ✗ | ✗ | ✗ | 1.5% | 111,101 |
|  | ✓ | ✗ | ✓ | ✗ | ✗ | 3.2% | 235,318 |
|  | ✓ | ✗ | ✗ | ✗ | ✗ | 20.6% | 1,521,474 |
|  | ✗ | ✗ | ✗ | ✗ | ✗ | 24.4% | 1,799,495 |
| NOA1 | ✓ | ✓ | ✓ | ✓ | ✓ | 28.4% | 1,606,970 |
|  | ✓ | ✓ | ✓ | ✗ | ✗ | 28.2% | 1,597,480 |
|  | ✓ | ✓ | ✗ | ✓ | ✗ | 0.0% | 1,586 |
|  | ✓ | ✓ | ✗ | ✗ | ✗ | 0.7% | 42,423 |
|  | ✓ | ✗ | ✓ | ✗ | ✗ | 4.0% | 228,219 |
|  | ✓ | ✗ | ✗ | ✗ | ✗ | 14.2% | 803,268 |
|  | ✗ | ✗ | ✗ | ✗ | ✗ | 24.5% | 1,385,299 |
| NOA2 | ✓ | ✓ | ✓ | ✓ | ✓ | 28.0% | 1,321,345 |
|  | ✓ | ✓ | ✓ | ✗ | ✗ | 28.4% | 1,337,904 |
|  | ✓ | ✓ | ✗ | ✓ | ✗ | 0.1% | 4,581 |
|  | ✓ | ✓ | ✗ | ✗ | ✗ | 2.1% | 99,454 |
|  | ✓ | ✗ | ✓ | ✗ | ✗ | 2.0% | 93,328 |
|  | ✓ | ✗ | ✗ | ✗ | ✗ | 15.3% | 720,629 |
|  | ✗ | ✗ | ✗ | ✗ | ✗ | 24.1% | 1,136,768 |

**Figure S1:**
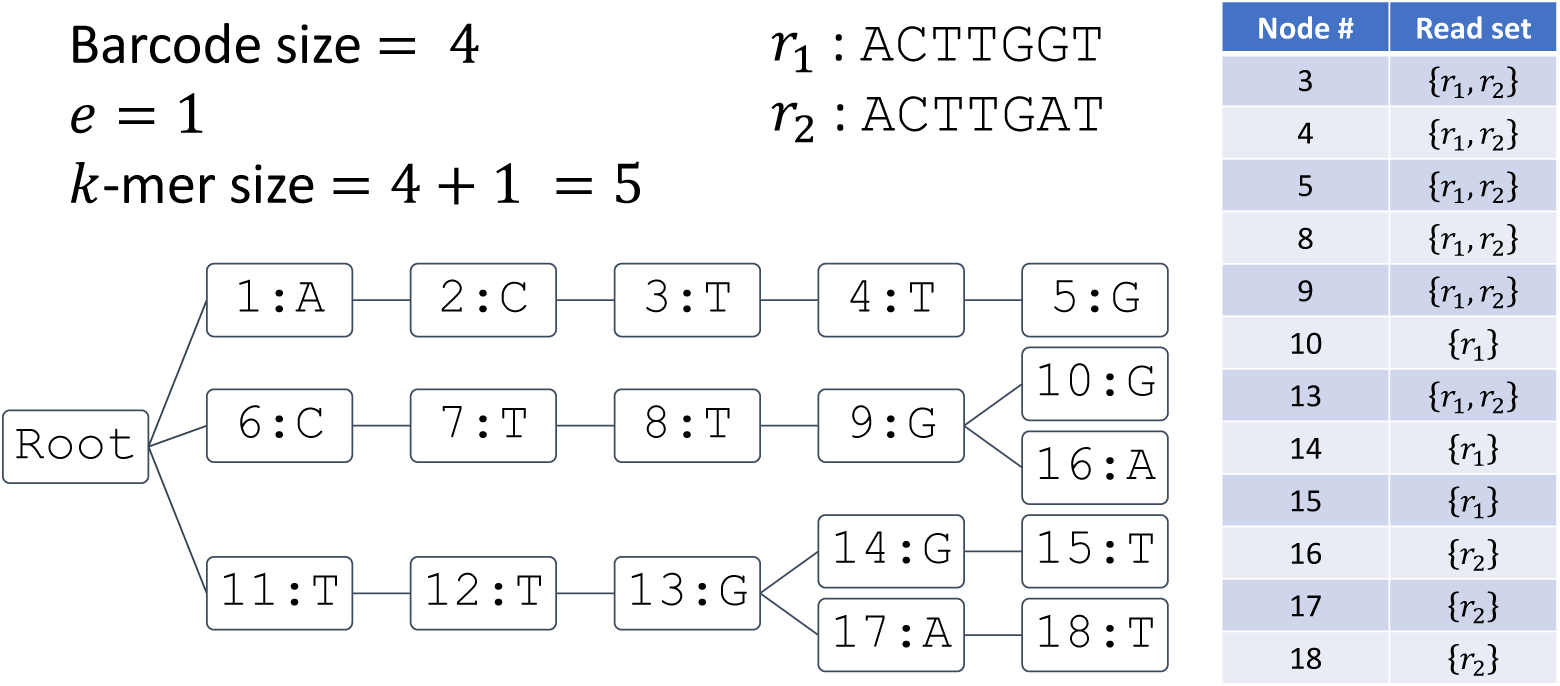
Illustrative example of trie construction in scTagger. Assuming barcode size of 4 and maximum allowed error *e* = 1, the *k*-mer size is 5. The *k*-mers of the two LR barcode segments, *r*_1_ and *r*_2_, are inserted into the trie. The nodes IDs correspond to the order by which they were inserted into the trie. Nodes at layers *k*−*e* = to *k*+*e* = (i.e. the deepest three layers in the example) contain nodes that barcode alignment can successfully terminate within the allowed error. A map with these nodes as keys is maintained. Each node maps to a set of LR segments from which we extracted *k*-mers that threaded through the node at insertion time.

### Algorithm 1 Automatic detection of the ranges of the adapter alignments on the LRs

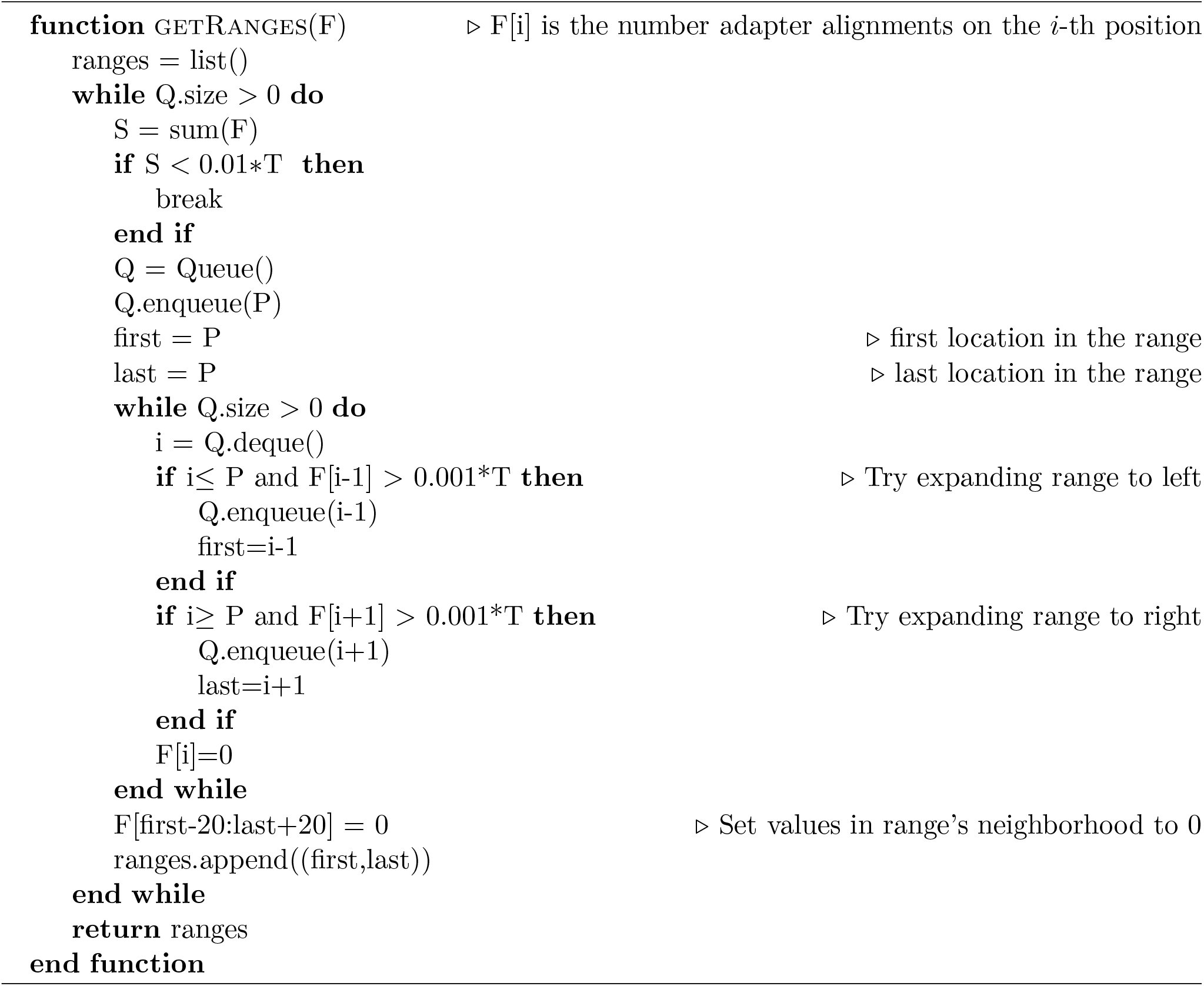

### Algorithm 2 Depth-first search in the trie

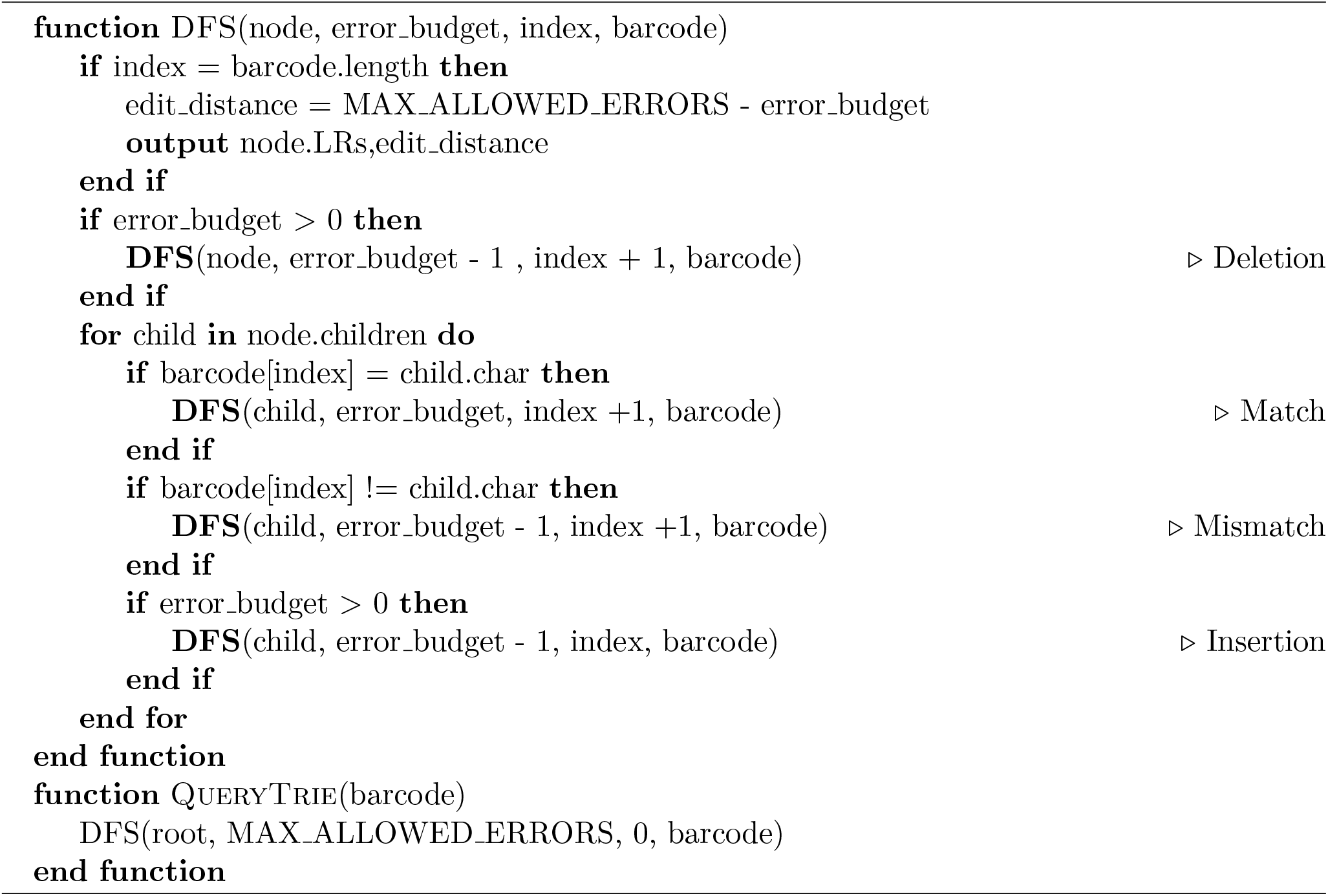

**Figure S2:**
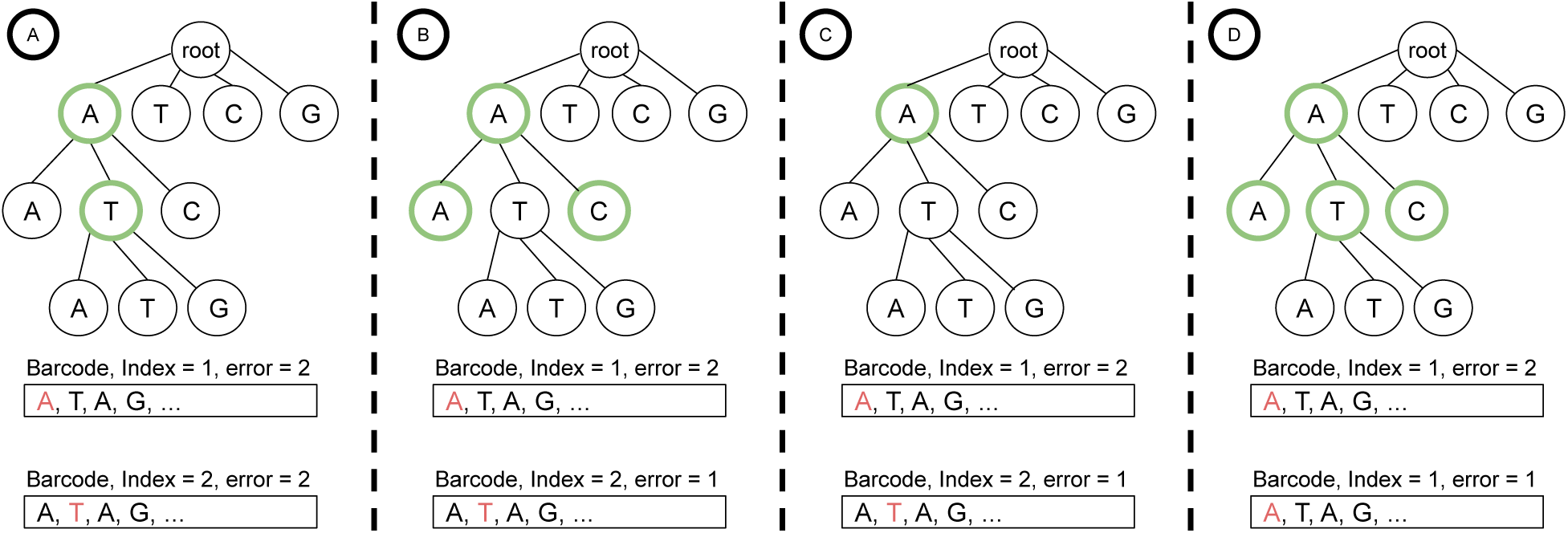
The different sub-queries of the search at a given node: A) Matching: If the a child node’s character matches the next barcode character, we explore this branch while incrementing the barcode index. B) Mismatch: If the a child node’s character does not match the next barcode character, we explore this branch while incrementing the barcode index and the total error incurred. C)Deletion: If a character is deleted from the long-read segment, we skip that corresponding barcode index by incrementing the index while stay on the same node in the trie and increment the total error incurred. D) Insertion: If a character is inserted in the long-read segment, we skip that node and to its children without increasing the barcode index while incrementing the total error incurred.

### Theorem S1.1.

*The time complexity for querying the trie in the matching stage of scTagger is O*(*Mϵ*^*e*^(*L* + *e*)^*e*+1^).

*Proof*. First, to prove the time complexity of querying the trie, we need to propose an upper-bound *U* on the number of possible strings with edit distances ≤ *e* from a barcode query assuming an alphabet with size *ϵ*. We claim that *U* = (2*ϵ* + 1)^*e*^ × (*L* + *e*)^*e*^. The length of the longest string with at most *e* edits is *L* + *e*. Therefore, the length of a string that we can still edit is at most *L* + *e* − 1. This is because the starting barcode has *L* characters and there could have been no more than *e*− 1 insertions applied before the current edit is applied. Keeping this in mind, we have three types of edits:

1. **Substitution**. We need to select a position to alter and then select another alphabet character to replace it. Since the string length is at most *L* + *e* − 1 before applying an edit, there are at most (*ϵ* − 1) × (*L* + *e* − 1) possible strings as a result of a substitution.
2. **Deletion**. We need to select a single position to delete. Therefore, we have at most *L*+*e* − 1 possible strings as a result of a deletion.
3. **Insertion**. We need to select a position and a character from the alphabet to insert at that position. Therefore, we have at most *ϵ* × (*L* + *e*) possible strings as a result of an insertion.
4. **No edit**. Additionally, we can decide not to use an edit, keeping the string unchanged. Obviously, there is 1 possible string resulting from this operation. This accounts for error distances strictly less than *e*.

Each of these options can be taken at most *e* times. Adding all this together (simplification: *L*+*e*−1 is upper bounded by *L* + *e*) gives us an upper bound of *U* = (2*ϵ* + 1)^*e*^ × (*L* + *e*)^*e*^, which proves the claim.

Finally, the cost of finding each of these possible strings in the trie is equal to the height of the trie tree, *L* + *e*. In total, we have the (2*ϵ* + 1)^*e*^ × (*L* + *e*)^*e*+1^ as the cost for each query barcode. And since we have *M* query strings, the total time complexity for the query stage of scTagger is *O*(*Mϵ*^*e*^(*L* + *e*)^*e*+1^).

## Notes

https://github.com/vpc-ccg/scTagger

### Competing Interest Statement

The authors have declared no competing interest.

